## Supplementary Notes 1-10 for "The DNA virome varies with human genes and environments"

### Supplementary Note 1 – Evidence supporting viral origin of sequenced reads

Several lines of evidence suggested that the reads aligned to viral genomes that passed our filtering pipeline (**Methods**) were of true viral origin:

1) For each virus, reads attributed to the virus covered the viral reference genome reasonably evenly (aggregating all reads observed to align to the virus across all individuals; **Extended Data Fig. 2**). Most reads attributed to a virus did not arise from any localized spike in read coverage, which would be a diagnostic sign of reads arising from mapping artifacts rather than from viral genomic DNA.
2) The presence of viral reads associated strongly with seropositivity for viral antigens measured in UKB (EBV: OR = 21.5, $p = 5.8\times10^{-28}$; HHV-7: OR = 2.5, $p = 9.7\times10^{-6}$; CMV: OR = 6.5, $p = 0.053$). For EBV, we directly estimated the false positive rate (that is, EBV alignments in individuals without evidence of prior EBV infection) using high-quality EBV serostatus data available in UKB. Among seronegative individuals, only 4 out of 494 individuals (0.8%) had EBV reads, and moreover, 2 of these individuals had three read pairs that mapped to EBV, suggesting that these reads were correctly attributed to EBV and that these 2 individuals had false negative serostatus. In contrast, 1,323 out of 8,857 seropositive individuals (14.9%) had EBV reads. This suggests that the false positive rate was well-controlled at roughly ~3% (= 2 / 492 / (1,323 / 8,857)).
3) For HHV-7 and HHV-6B, which we observed in ≥90% of saliva DNA samples from adults (for HHV-7) or children age 5–9 (for HHV-6B) (**Fig. 2a**), viral reads were observed in only a small minority of the youngest SPARK participants (infants) (**Extended Data Fig. 1d**). This observation is consistent with infants being unlikely to have been infected and indicates that the false positive rate of viral reads was well-controlled. Similarly, MCPyV, which was observed in ~10–20% of young children in SPARK, was not observed in any infants under 5 months of age.
4) The prevalence of viral reads tracked with lymphocyte percentage (measured by hematological assays in UKB) for herpesviruses known to latently reside in lymphocytes (EBV in B cells and HHV-6B and HHV-7 in T cells[1]) but not for anelloviruses, which infect a broader range of cell types[2] (**Extended Data Fig. 1c**).

1. Mori, Y. & Yamanishi, K. HHV-6A, 6B, and 7: pathogenesis, host response, and clinical disease. in *Human Herpesviruses: Biology, Therapy, and Immunoprophylaxis* (eds Arvin, A. et al.) (Cambridge University Press, Cambridge, 2007).

2. Spandole, S., Cimponeriu, D., Berca, L. M. & Mihăescu, G. Human anelloviruses: an update of molecular, epidemiological and clinical aspects. *Arch. Virol.* **160**, 893–908 (2015).

#### Supplementary Note 2 – Robustness of results to viral alignment mapping quality

The analyses presented in this work used alignments to viral genomes with a mapping quality threshold of 5 (i.e., MAPQ≥5) to maximize recovery of reads with some variation from viral reference genomes (**Methods**). To evaluate sensitivity of our conclusions to mapping quality, we repeated the analyses of UK Biobank using a more stringent filter (MAPQ≥20) and found that this affected very few reads (and very few individuals). For example, for the 6 most prevalent viruses, the affected individuals (i.e., those who had alignments with MAPQ between 5 and 19) represented only a modest fraction of the number of individuals who we had determined to be viral DNA-positive based on the MAPQ≥5 threshold: 0.5% of individuals with alignments to the EBV genome (367/72157), 1.8% for HHV-7 (783/42987),1.8% for HHV-6B (106/5675), 2.5% for TTV (TUS01) (143/5582), 0.3% for TTV (HD14a) (22/6264), and 7.9% for TTV (VT416) (171/1990). The relatively larger fractions of affected individuals for two anelloviruses (TUS01 and VT416) probably reflect the higher sequence diversity of anelloviruses, which results in more low-quality alignments due to mutations relative to the viral reference genomes.

We further verified that the choice of mapping quality threshold had a negligible impact on genetic associations we identified with viral DNA load. For example, the top HLA associations for each of the above viruses (excluding HHV-6B, which lacks an association in the MHC region) were nearly unchanged upon switching from MAPQ≥5 to MAPQ≥20: the strongest EBV association changed from $p = 2.6 \times 10^{-670}$ to $9.2 \times 10^{-669}$; for HHV-7 $p = 6.9 \times 10^{-603}$ to $1.5 \times 10^{-608}$; for TTV (TUS01) $p = 2.0 \times 10^{-80}$ to $9.4 \times 10^{-81}$; for TTV (HD14a) $p = 5.8 \times 10^{-251}$ to $4.0 \times 10^{-252}$; for TTV (VT416) $p = 1.1 \times 10^{-16}$ to $1.1 \times 10^{-14}$.

#### Supplementary Note 3 – Robustness of temporal trends to potential confounders

We performed multiple lines of sensitivity analyses to test whether observed temporal patterns were robust to several variables that might directly bias results (for example, an individual's sex or age) or encode latent information about individuals, such as assessment center. We included age, age-squared, sex, assessment center, and top genetic principal components as covariates (**Methods**) in models used to generate primary results (**Fig. 2e,f** and **Extended Data Fig. 3k-n**). Including day-of-week (encoded as indicator variables) had no effect on either EBV or HHV-7 ($p>0.05$ for all indicator variables) when included as a covariate in both the circadian and seasonal analyses. No assessment center had an outlier rate of EBV or HHV-7 positivity: EBV prevalence ranged from –0.080 to +0.042 s.d. and HHV-7 prevalence ranged from –0.047 to +0.11 s.d., so we concluded that no center needed to be excluded from analyses. Circadian and seasonal patterns were broadly consistent within each assessment center (different colored lines in **Extended Data Fig. 3e-h**). Likewise, circadian patterns were consistent within each season (**Extended Data Fig. 3i,j**). However, we cannot rule out the possibility that some of these temporal associations may be confounded by unmeasured visit-related factors. Independent replication in future cohorts will clarify interpretation of these results.

#### Supplementary Note 4 – Variation in viral DNA load across ancestries

EBV DNA was least commonly observed in European-ancestry donors and most commonly observed in blood samples from African-ancestry individuals, varying in prevalence by nearly 2-fold across ancestries in both UKB ($p$ = 3.2x10$^{-293}$ by regression, **Extended Data Fig. 4a**) and AoU ($p$ = 4.8×10$^{-1524}$, **Extended Data Fig. 4b**). Surprisingly, these ancestry effects were somewhat different in saliva samples, with individuals of East Asian and American ancestries exhibiting the highest EBV prevalence and abundance in saliva, consistently across AoU and SPARK ($p$ = 6.4×10$^{-184}$ and 2.4×10$^{-20}$ in AoU; $p$ = 2.6×10$^{-8}$ and 0.073 in SPARK, **Extended Data Fig. 4c-f**). In admixed AoU participants, EBV viral read prevalence in blood WGS tracked with proportions of African and American admixture ($p$ = 1.2×10$^{-763}$ and 1.1×10$^{-696}$, **Extended Data Fig. 4g,h**). Other viruses exhibited even more variation in blood viral DNA load across ancestries: HHV-7 viral read prevalence was 2–4 fold higher in African-ancestry individuals compared to other ancestries ($p$ = 3.2×10$^{-495}$ and 4.8×10$^{-1319}$, **Extended Data Fig. 4i,j**), and TTV viral read prevalence exhibited a 5–9-fold range, from 2.19% (2.15–2.23%) in European-ancestry UKB participants to 18.8% (16.8–20.8%) in participants of East Asian ancestry ($p$ = 1.6×10$^{-1318}$ and 4.5×10$^{-3011}$, **Extended Data Fig. 4k,l**).

To evaluate whether the large observed differences in viral DNA loads between genetic ancestries could be explained in part by socioeconomic correlates, we examined the relationship between viral DNA loads measured in UK Biobank blood samples and the following variables: Townsend deprivation index (a widely used measure of material deprivation in geographic areas), number of years spent in education, and smoking status. Although all of these variables associated significantly with DNA load of EBV and HHV-7, they all had a negligible impact on the effect size or significance of the effect of genetic ancestry when included as covariates in a model alongside standard covariates (age, age-squared, sex, assessment center). For example, in an analysis of individuals with either European or African genetic ancestry, African ancestry as a covariate in a model without Townsend deprivation index, educational years, or smoking status had an effect size of 0.24 s.d. [0.22,0.25] (p=4.6x10$^{-227}$) on EBV DNA load (i.e., inverse-normal transformed number of observed read pairs), whereas in a model with these covariates, African ancestry had an effect size of 0.23 s.d. [0.21,0.24] (p=4.9x10$^{-194}$). For HHV-7, the effect size changed from 0.32 s.d. [0.30,0.33] (p=3.5x10$^{-512}$) to 0.32 s.d. [0.31,0.33] (p=1.4x10$^{-491}$). Likewise, in an analysis of individuals with either European or South Asian genetic ancestry, the effect size for South Asian ancestry on EBV DNA load changed from 0.17 s.d. [0.15,0.18] (p=1.7x10$^{-140}$) to 0.17 s.d. [0.15,0.18] (p=1.0x10$^{-128}$). For HHV-7, the effect size changed from -0.090 s.d. [-0.10,-0.079] (p=1.9x10$^{-55}$) to -0.087 s.d. [-0.099,-0.076] (p=3.7x10$^{-48}$). Overall, these results suggest a minimal contribution of socioeconomic factors to the differences in viral DNA loads across genetic ancestries.

#### Supplementary Note 5 – Consistency of genetic associations with viral DNA detection across viral genomes

Associations of human genetic variants with WGS-derived viral DNA load phenotypes could in theory reflect either true genetic associations with viral DNA load or misalignment of human DNA sequences to a viral genome. To evaluate the latter possibility, we examined whether lead variants identified by our GWAS of EBV and HHV-7 viral DNA load consistently associated with the presence of reads aligned to the left half of the viral genome and with reads aligned to the right half. If an association of a human genetic variant with viral DNA load were an artifact of misalignment, it should be driven by read alignments to a specific region of the virus's genome (at which human sequences misalign) and not associate with the presence of reads aligned to the half of the viral genome not containing this region. We found that all of the lead variants (except one false positive flagged based on inconsistent effect directions in UKB and AoU; see **Methods**) had comparable associations with reads from the right and left halves of each viral genome (**Extended Data Fig. 7a,b**), suggesting that these associations are unlikely to be due to human allele-specific misalignment to viral genomes.

### Supplementary Note 6 – Telomeric integration sites of HHV-6A and HHV-6B

Genome-wide association analyses of chromosomally integrated (endogenous) HHV-6A and HHV-6B found strong associations with peritelomeric variation (**Extended Data Fig. 7j,k**). As in previous work[24,25], these associations presumably reflect haplotypes near the telomere that contain the inherited integration, rather than variants that influence integration itself. These associations indicate that at least seven HHV-6B integration events in seven telomeres (7p, 9p, 9q, 11p, 17p, 19q, and 21q; **Extended Data Fig. 7k**) have generated eHHV-6B haplotypes in European-ancestry populations, and at least three HHV-6A integration events in three telomeres (17p, 18q, and 19q; **Extended Data Fig. 7j**) have generated eHHV-6A haplotypes in European-ancestry populations. This is notable compared to the single common 22q integration seen in 72% of Asian individuals with inherited HHV-6[1]. In fact, we observed significantly higher rates of endogenous HHV-6A and HHV-6B in individuals of European ancestry (0.33% and 1.09%) than in those of African (0.09% and 0.23%, $p = 4.4 \times 10^{-5}$ and $4.1 \times 10^{-18}$ by Fisher's exact test), East Asian (0.07% and 0.27%, p = 0.10 and 0.00062), and South Asian (0.11% and 0.12%, $p = 5.6 \times 10^{-5}$ and $2.4 \times 10^{-30}$) ancestry in UK Biobank.

1. Liu, X. *et al.* Endogenization and excision of human herpesvirus 6 in human genomes. *PLOS Genetics* **16**, e1008915 (2020).

#### Supplementary Note 7 – Associations of quantitative antibody responses to viral antigens with viral DNA load and with MHC variation

Quantitative antibody responses measured in a subset of ~10,000 UK Biobank participants have previously been observed to associate with human genetic variation in the MHC region[1,2] and might be expected to correlate with viral DNA load. Surprisingly, we found minimal concordance in the MHC region between genetic associations with EBV DNA load and associations with quantitative antibody responses to EBV antigen EBNA-1 (**Extended Data Fig. 8a**). Interestingly, antibody responses to EBNA-1 had a stronger association with the multiple sclerosis risk allele[3], *DRB1*15:01* ($p = 1.1 \times 10^{-31}$), than *DRB1*04:04* ($p = 2.4 \times 10^{-6}$). Associations with antibody response against other EBV antigens (ZEBRA and VCA-p18) showed more concordance with genetic effects on EBV DNA load at the MHC (**Extended Data Fig. 8c,d**). Accordingly, the number of WGS read pairs mapping to the EBV genome was associated with quantitative antibody responses against ZEBRA ($p = 5.2 \times 10^{-15}$) and VCA-p18 ($p = 8.1 \times 10^{-25}$), more weakly with the response against EA-D ($p = 6.0 \times 10^{-4}$), but not with the response against EBNA-1 ($p = 0.053$, **Extended Data Fig. 8f-i**). Likewise, we found poor concordance between the genetic associations to quantitative antibody response against HHV-7 U14 and HHV-7 DNA load (**Extended Data Fig. 8e**) as well as a weak association between the two phenotypes ($p = 8.2 \times 10^{-4}$; **Extended Data Fig. 8j**). Overall, this suggests that some quantitative antibody responses, such as those against VCA-p18 and ZEBRA, are likely partially influenced by viral DNA load, whereas others, such as those against EBNA-1, EA-D, and U14, appear to be largely independent and might reflect variation in immunological response.

1. Kachuri, L. *et al.* The landscape of host genetic factors involved in immune response to common viral infections. *Genome Medicine* **12**, 93 (2020).

2. Mentzer, A. J. *et al.* Identification of host–pathogen-disease relationships using a scalable multiplex serology platform in UK Biobank. *Nat Commun* **13**, 1818 (2022).

3. Multiple Sclerosis Genomic Map implicates peripheral immune cells & microglia in susceptibility. *Science* **365**, eaav7188 (2019).

#### Supplementary Note 8 – Complex effects of MHC variation on viral DNA load

The strong effects of genetics and age on viral DNA load led us to wonder how genetic effects on viral DNA load might themselves vary across donors. For both EBV and HHV-7, the effect size of the lead variant in the MHC region (*HLA-DRB1*04:04* for EBV) was greater in age ranges with higher mean viral DNA load (**Extended Data Fig. 9a,b**). Variants at MHC appeared to generally act multiplicatively (rather than additively) with the effects of genetic variation elsewhere in the human genome (**Extended Data Fig. 9c,d**), deviating from the standard model of additive polygenic effects. The strong effects from the MHC region also exhibited complex interactions with one another. Variants in the MHC region generally associated with effects on EBV DNA load that were consistent among *DRB1*04:04* carriers and non-carriers, but we observed several exceptions potentially reflecting interactions between the set of presented peptides presented by *DRB1*04:04* and other alleles in generating an effective immune response. (**Extended Data Fig. 9e**). The *DRB1* T210 allele associated with decreased EBV DNA load in noncarriers of *DRB1*04:04* but with increased EBV DNA load in carriers (**Extended Data Fig. 9f**), and the *DQA1* E198 allele exhibited a similar reversal (**Extended Data Fig. 9g**). The *DPB1* V105 allele also appeared to display strongly non-additive interactions with *DRB1*04:04* (**Extended Data Fig. 9h**).

#### Supplementary Note 9 – Association of Duffy-null genotype with viral DNA load and neutropenia

To identify ancestry-enriched genetic variants that might contribute to viral load differences between ancestries, we conducted GWAS for EBV and HHV-7 DNA load in blood restricted to individuals of African ancestry in AoU (n=77,573, **Methods**). These two GWAS primarily re-identified variants that we had observed to associate with viral DNA load in our larger GWAS of European-ancestry individuals, with one notable exception: variants within a wide ~10Mb region on chromosome 1q associated with both EBV and HHV-7 DNA load, characteristic of admixture-induced linkage disequilibrium involving a causal variant with divergent allele frequencies in the recently-admixed parent ancestries[1] (**Extended Data Fig. 10a,b**).

The top associated variant (rs2814778) is the known African-specific (AF = 0.82 in AoU AFR vs. 0.01 in AoU EUR) variant in the promoter of the *ACKR1* gene that causes the recessive Duffy-null phenotype by disrupting a GATA-binding site, preventing surface expression of the Duffy antigen on red blood cells[2]. The Duffy-null phenotype most notably confers malaria resistance[3]. In the AoU AFR cohort, rs2814778-CC (Duffy-null; 67.8% of AoU AFR participants) associated with 2.7% [2.0%,3.5%] and 5.1% [4.4%,5.8%] higher prevalence of EBV and HHV-7 DNA positivity, respectively (linear regression of binary positivity on rs2814778-CC vs. rs2814778-CT/TT, with age, age squared, sequencing center, and 16 genetic PCs as covariates). To ensure that this association was not confounded by variation in AoU participants' genome-wide AFR vs. EUR ancestry proportions (which is modestly correlated with rs2814778), we confirmed that it was robust to restricting to a subset of the AoU AFR cohort within a narrow range of the spectrum of genome-wide AFR ancestry fractions (specifically, individuals with 70-90% African ancestry and >90% AFR+EUR ancestry; n=51,052, **Extended Data Fig. 10c**). Within this sub-cohort, the mean African ancestry proportion was only 2% higher in Duffy-null individuals (83.5%) compared to non-Duffy-null individuals (81.2%), and the effect sizes of rs2814778-CC on viral DNA positivity were essentially unchanged (3.2% [2.3%, 4.1%] for EBV, 5.2% [4.4%, 6.1%] for HHV-7.

To quantify the proportions of the African vs. European differences in prevalence of EBV and HHV-7 DNA positivity explained by the Duffy-null genotype, we computed the ratio of the prevalence difference among Duffy-null AFR vs. non-Duffy-null AFR participants to the prevalence difference among Duffy-null AFR vs. EUR participants in AoU, obtaining estimates of 28% [21%, 35%] for EBV and 45% [38%, 52%] for HHV-7. This calculation treats the Duffy-null AFR population as a proxy for non-admixed African-ancestry populations (in which the Duffy-null allele is close to fixation; e.g., AF = 0.999 in 1000 Genomes non-admixed AFR populations). An alternative calculation treating all 51,052 individuals in the sub-cohort above (**Extended Data Fig. 10c**) as the AFR reference population (i.e., computing the ratio of the prevalence difference among AFR vs. non-Duffy-null AFR participants to the prevalence difference among AFR vs. EUR participants in AoU) gives estimates of 21% [14%, 28%] for EBV and 37% [30%, 45%] for HHV-7.

One caution we note in interpreting the associations of Duffy-null genotype with EBV and HHV-7 DNA positivity is that aside from malaria resistance, the other major phenotype associated with

Duffy-null genotype is reduced neutrophil count[4,5]. This could contribute to an increase in measured viral DNA load (based on positivity or counts of viral DNA in blood-derived WGS data) by increasing lymphocyte percentage (**Extended Data Fig. 1c**), as lymphocytes and neutrophils compose the majority of white blood cells from which blood-derived DNA is extracted, and EBV and HHV-7 reside latently in episomes within lymphocytes. We confirmed that in UKB African-ancestry individuals (n=7,392), the Duffy-null genotype associated strongly ($p$=4.9x10$^{-115}$) with an 8.2% increase in lymphocyte percentage in a model including standard covariates (age, age-squared, sex, assessment center). Similarly, in AoU African-ancestry individuals for whom we could generate a lymphocyte percentage phenotype (n=28,051; **Methods**), the Duffy-null genotype associated strongly ($p$=7.7×10$^{-440}$) with 6.3% higher lymphocyte percentage in a model including standard covariates (age, age-squared, sex, site, ancestry PCs). However, these increases in lymphocyte percentage would be expected to produce only ~1–2% increases in rates of observed EBV and HHV-7 DNA positivity (based on our analyses in UKB; **Extended Data Fig. 1c**), suggesting that the Duffy-null effect on neutrophil count was unlikely to fully explain its effect on ancestry differences in viral DNA prevalence.

To evaluate this question more directly, we explored the extent to which adjusting for lymphocyte percentage affected associations of Duffy-null genotype with viral DNA load measurements. To maximize statistical power, we analyzed inverse normal transformed EBV and HHV-7 abundance phenotypes in AoU blood samples from African-ancestry individuals (n=16,451), including standard covariates (age, age-squared, sex, site, ancestry PCs). The effect size of the Duffy-null genotype on EBV DNA load decreased from 0.062 s.d. [0.035,0.089] ($p$=7.2x10$^{-6}$) to 0.044 s.d. [0.016,0.072] ($p$=0.0022) upon including lymphocyte percentage as a covariate. Likewise, for HHV-7 DNA load, the effect size decreased from 0.10 s.d. [0.074,0.13] ($p$=1.2x10$^{-13}$) to 0.077 s.d. [0.049,0.10] ($p$=4.4x10$^{-8}$). These analyses suggest that the Duffy-null effect on viral DNA load is not mediated solely by its effect on reducing neutrophil abundance (thereby shifting blood cell composition). However, a limitation of this analysis is that the lymphocyte percentage phenotype evaluated here (obtained from an AoU participant's electronic health records) is an imperfect proxy for the lymphocyte percentage of the blood sample from which DNA was extracted (for example, due to temporal variation in lymphocyte percentage or simply measurement error).

An orthogonal way to evaluate whether the Duffy-null phenotype influences viral DNA load independently of its effect on neutrophil abundance in blood is to test for an effect in saliva. As viral alignments in saliva are much more abundant and likely primarily derived from viral particles, their abundance should be unaffected by the Duffy-null effect on blood composition. In AoU saliva samples, HHV-7 abundance was significantly higher in Duffy-null individuals ($p$=0.027 in a linear model with standard covariates; **Extended Data Fig. 10d**). We did not observe a significant effect for EBV ($p$=0.84).

Overall, these lines of evidence lead us to conclude that the Duffy-null phenotype explains a substantial fraction of the African vs. European differences in EBV and HHV-7 DNA load, and that these effects—particularly for HHV-7—are unlikely to be fully explained by neutropenia in Duffy-null individuals.

#### Supplementary Note 10 – Limitations of present study

While biobank WGS data sets offer the opportunity to profile DNA viromes at an unprecedented scale (requiring only computational analyses of existing data), the individual-level viral DNA load phenotypes that we generated were noisy, stochastic observations, often of only a single viral DNA fragment present in a sequenced blood sample. These analyses will need to be revisited in future studies of even larger population biobank cohorts that increase the number of genetic instruments available for Mendelian randomization, which will improve the resolution and robustness of MR analyses. Saliva samples, in which we found viral DNA to be more abundant, allowed assessing DNA load for more viruses with greater precision, but most WGS data have been generated from blood. Studying the virome at higher resolution and in other body sites will require other data sources[1,2]. Additionally, although we could infer average trajectories of viral DNA load with age, time of day, and time of year using cross-sectional analyses, studying individual-level viral DNA load trajectories will require longitudinal sampling[3]. The analytical pipeline that we utilized also imposed limitations: future work could expand the set of viral genomes considered to include other common DNA viruses (e.g., adeno-associated viruses or parvovirus 4) and undertake further strain-level genetic association analyses, which we used here to fine-map complex genetic associations at HLA with EBV. Our analytical approach of aligning sequencing reads to reference genomes also reduces the ability to observe reads from viruses with more divergent genomes. One possible approach could be to use modern translated search alignment software such as DIAMOND[4] to better capture reads derived from such genomes. Finally, the cohorts we studied here, from the UK and United States, provided limited representation of viruses that are endemic in other parts of the world.

1. Kumata, R., Ito, J., Takahashi, K., Suzuki, T. & Sato, K. A tissue level atlas of the healthy human virome. *BMC Biol.* **18**, 55 (2020).

2. Pyöriä, L. *et al.* Unmasking the tissue-resident eukaryotic DNA virome in humans. *Nucleic Acids Res.* **51**, 3223–3239 (2023).

3. Khan, G., Miyashita, E. M., Yang, B., Babcock, G. J. & Thorley-Lawson, D. A. Is EBV persistence in vivo a model for B cell homeostasis? *Immunity* **5**, 173–179 (1996).

4. Buchfink, B., Reuter, K. & Drost, HG. Sensitive protein alignments at tree-of-life scale using DIAMOND. *Nat Methods* **18**, 366–368 (2021).
